# Asymmetrical adaptations to increases and decreases in environmental volatility

**DOI:** 10.1101/2021.07.30.454486

**Authors:** Jie Xu, Nicholas T. Van Dam, Yuejia Luo, André Aleman, Hui Ai, Pengfei Xu

**Author notes:** Corresponding author: Hui, Ai, Ph.D. Center for Brain Disorders and Cognitive Sciences, Shenzhen University, Shenzhen, China, Pengfei Xu, Ph.D. Faculty of Psychology, Beijing Normal University, Beijing, China.

## Abstract

Humans adapt their learning strategies to changing environments by estimating the volatility of the reinforcement conditions. Here, we examine how volatility affects learning and the underlying functional brain organizations using a probabilistic reward reversal learning task. We found that the order of conditions was critically important; participants adjusted learning rate going from volatile to stable, but not from stable to volatile, environments. Subjective volatility of the environment was encoded in the striatal reward system and its dynamic connections with the prefrontal control system. Flexibility, which captures the dynamic changes of network modularity in the brain, was higher in the environmental transition from volatile to stable than from stable to volatile. These findings suggest that behavioral adaptations and dynamic brain organizations in transitions between stable and volatile environments are asymmetric, providing critical insights into the way that people learn under uncertainty.

## Introduction

Learning in an uncertain environment requires flexibility, appropriately adjusting perceived action-outcome associations. Individuals must adjust quickly in dynamic environments to update prior estimation of the association between action and outcome. In stable environments, people relaxedly fine-tune strategies to maintain beliefs about unchanging associations. Flexibly adjusting learning strategies between different environments is critical to optimal performance *(1, 2)*. However, adaptations to different environments largely rely on the order in which people experience changes *(3)*. People may adapt more quickly in one direction (e.g., moving from a more to less effortful context relative to its opposite). Transfer effects, for example, have historically been shown to favor the difficult to easy direction *(4)*.

Uncertainty is an inherent structure of the environment, the estimates of which could be used to characterize organismal adaptability *(5)*. Most theoretical propositions of uncertainty are focused on three key types of uncertainty: irreducible uncertainty, estimation uncertainty, and unexpected uncertainty *(5-7)*. Irreducible uncertainty, the first-order uncertainty, is represented by risk, wherein the probability of an option-outcome association is known but the outcome of reward or punishment remains uncertain *(8)*. Second-order (estimation) uncertainty reflects ambiguity, wherein the probability of a stimulus being associated with a given outcome is unknown and needs to be estimated *(8)*. Unexpected (or third-order) uncertainty is volatility, the frequency at which the association between the stimulus and outcome varies dynamically. Irreducible (first-order) and estimation (second-order) uncertainty constitute *expected or known uncertainty (5, 7)*. In responses to *expected uncertainty*, individuals need more observations to estimate the state of the environment but have to ignore or discount specific surprise outcomes, whereas decisions in *unexpected uncertainty* or volatility rely on the most recent observations given that the association between options and outcomes are likely changing. To adapt to a complex environment, it is necessary to identify and estimate the expected and unexpected uncertainty leading to a surprise event, which may reflect a major change in action-outcome associations. Recent studies suggest that individuals make use of uncertainty to guide their decisions during the learning process *(6, 9, 10)*. Difficulties in estimating environmental uncertainty may contribute to maladaptive functions in internalizing psychological disorders (e.g., anxiety, depression) *(11, 12)*. Although previous studies have shown different learning rates between stable and volatile environments *(1, 2, 11)*, adaptation processes and brain function underlying directional changes in volatility (i.e., the transition from stable to volatile environment vs. from volatile to stable environment) remain unclear.

Adaptation to uncertainty is underpinned by dynamic and distributed brain networks. The anterior cingulate cortex (ACC), a part of the salience network which detects error and conflict, fluctuates co-incidentally with estimated volatility *(1, 2, 9, 13)*. Key regions of the default mode network, such as the posterior cingulate cortex (PCC), have also shown to be negatively correlated with unexpected uncertainty *(8, 10, 13)*. The orbitofrontal cortex (OFC) and caudate, parts of the reward network, have shown increased activity during changing learning and reward probabilities *(9, 14-17)*. One recent study has also shown that surprise and uncertainty during learning are dynamically encoded by the frontoparietal control network, which is linked to appropriate behavioral adaptation *(18)*. Network flexibility, measured by the association of nodes to modules of brain networks, has also been shown to positively predict new learning *(19)*. Therefore, the combination of quantitative behavioral models and brain network models, have considerable promise for understanding human learning *(20)*.

In this study, we examined how the human brain adapts to transitions in environmental volatility (i.e., the transition from volatile to stable environments in comparison to transitions from stable to volatile environments). Given the historical transfer effect (easier going from hard to easy than vice versa *(4)*), we predicted more effortful adaptation in the direction from volatile to stable environments than the other way around. Dynamic organization of the brain networks underlying the transfer effect associated with environmental volatility were tested using dynamic brain network analyses on functional magnetic resonance imaging (fMRI) data.

## Results

Thirty-seven participants were asked to complete an adjusted probabilistic reward reversal learning task while undergoing functional magnetic resonance imaging (fMRI). On each trial, participants had to choose one of two options with specific reward probabilities (Fig. 1a). In the stable block, two options are consistently associated with reward probabilities of 75% and 25%, respectively (Fig. 1b). In the volatile block, the reward probabilities associated with the options changed between high (80%) and low (20%) every 20 trials (Fig. 1c). To perform optimally, participants had to estimate the reward probabilities of two options from the outcome of previous trials. Option selection and reaction time were recorded. Brain activity was also measured. Three participants were excluded from all analyses as they missed more than 10 of the 180 trials during the primary task (5.6%).

**Fig. 1.**
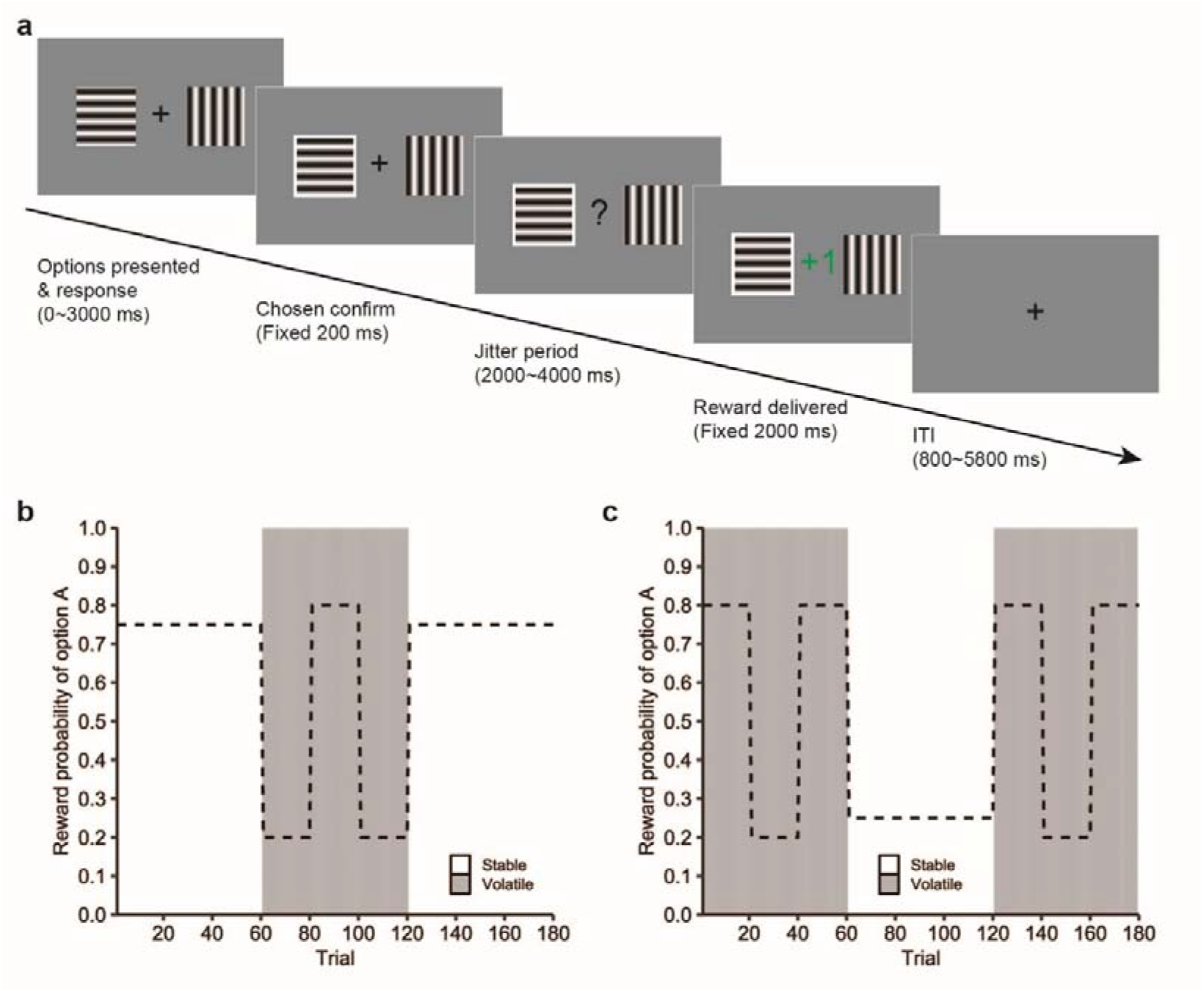
Task design. **a) Experimental procedure**. Participants were required to choose one of the two options with either horizontal or vertical gratings to maximize reward. A response cue indicated which option had been selected. A variable jitter was followed by reward presentation. In the outcome phase, the reward (green “+ 1”) or the no reward (red “+ 0”) was presented for two seconds. **b) Reward probabilities across the course in the stable-volatile-stable task**. This task consisted of three blocks (stable-volatile-stable). In the stable block, one option was linked to a reward with 75% probability while the other one would be followed by a reward with 25% probability. In the volatile block, the reward probabilities of the two options would switch between 20% and 80% every 20 trials. **c) Reward probabilities across the course in the volatile-stable-volatile task**. This task consisted of three blocks (volatile-stable-volatile). The reward probability of options was same with the one in the stable-volatile-stable task.

### Behavioral Results

Learning rate (α) reflects how participants’ choice at a given time was influenced by recent previous outcomes. A high learning rate means that current choice is strongly guided by recent outcomes and reflects rapidly changing stimulus-outcome associations. High learning rates are more suitable for volatile environments. In contrast, a low learning rate means that a surprising outcome has little effect on the subsequent choice. Low learning rates are more suitable to stable environments, reflecting that people may not change their selections. We estimated learning rates by fitting a Rescorla-Wagner (RW) learning model to their choices in the three blocks ^2, 11^. We also used a Bayesian learning model, which estimates the volatility of the task ^2, 11^. Model comparison indicated that the RW model using Grid search provided the best model-to-data fit (RW Grid ML: 115.61 ± 38.57, RW Grid EV: 117.58 ± 37.79, RW fMIN: 122.96 ± 37.88, Bayesian: 260.02 ± 53.82, *F*_(3, 99)_ = 139.5, *p* < 0.001; Fig. 2a). There was no difference between the ML estimate and EV estimate *(p* = 0.931). The RW model with Grid Search also generated the largest correlation between modeled probability and participants’ choices (RW Grid ML: 0.801 ± 0.110, RW Grid EV: 0.801 ± 0.108, RW fMIN: 0.771 ± 0.104, Bayesian: 0.736 ± 0.699, *F*_(3, 99)_ = 10.615, *p* < 0.001; Fig. 2b). Therefore, the Bayesian model was only used to estimate environmental volatility.

**Fig. 2.**
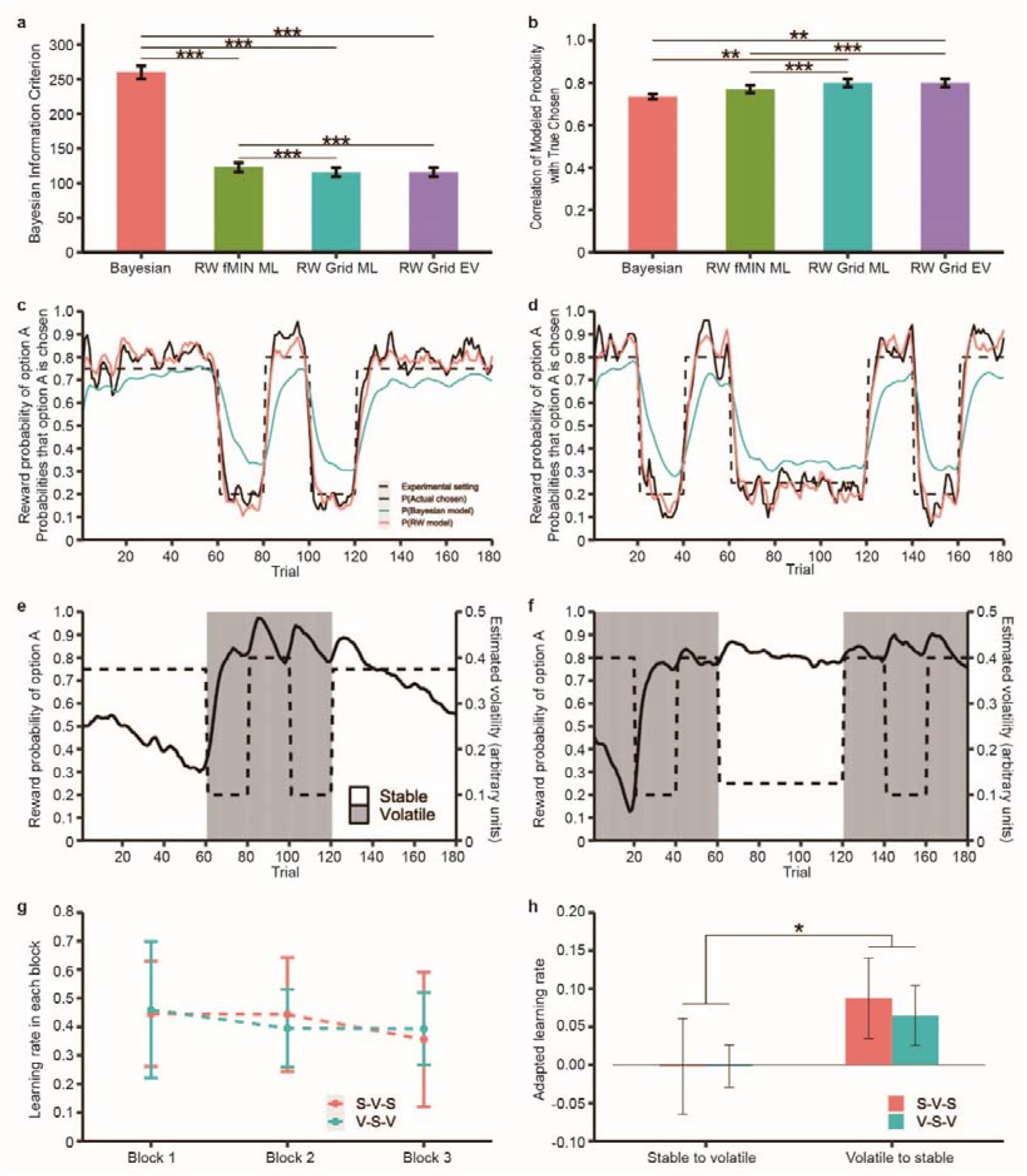
Behavioral results. **a) Model comparison by Bayesian information criterion (BIC).** The RW Grid model shows the lowest BIC among all models. Error bars represent the standard error. **b) Model comparison by the correlation of modeled probability with participants’ choices**. The RW Grid models showed the largest correlations among all models. Error bars represent the standard error. **c d) Learning curves illustrating participants’ choices and estimates of model during the c) Stable-Volatile-Stable task** and **d) Volatile-Stable-Volatile task**. The black dashed line represents the experiment setting of the probability of the highly rewarded option. The black solid line represents the participant’s choices. The salmon solid line represents the Rescorla-Wagner model prediction. The dark turquoise solid line represents the Bayesian model prediction. To illustrate participants’ choices and model predictions, the variation across the participants and trials has been reduced by smoothing using a running average of four trials. **f) Environmental volatility estimated by the Bayesian model across the whole experiment. e) Stable-Volatile-Stable task**. In the stable-volatile-stable task, estimated volatility (black solid lines, righthand axes) decreased gradually in the stable states and increased suddenly after each reversal. **f) Volatile-Stable-Volatile task**. In the volatile-stable-volatile task, after the first volatile state, estimated volatility (black solid lines, righthand axes) decreased slowly in the stable state. The underlying reward probabilities of the task are presented with black dashed lines (lefthand axes). **g h) Learning rates in each state. g) Learning rates fitted by the Rescorla-Wagner model to choices in each block of each task**. Salmon dashed line for stable-volatile-stable task. Dark turquoise dashed line for volatile-stable-volatile task. Dots represented the mean of participants’ learning rates. Error bars represented the standard deviation of participants’ learning rates in each block. **h) The adapted learning rates were significantly higher from volatile to stable state than it from stable to volatile state, regardless of the order in which the two blocks were first completed**. Error bars represent the standard error.

To understand how participants’ behaviors and mental representations changed over time, we examined participants’ cumulative true responses and modeled choices. In general, participants could successfully capture changes in probability across the whole experiment. Compared to the Bayesian model, the RW model fitted behavioral data better in both the stable-volatile-stable task and the volatile-stable-volatile task (Fig. 2c, d). We also estimated the environmental volatility *ν*_*(i)*_ by using the Bayesian model across the whole task. The estimated volatility increased suddenly after reversal and decreased gradually in the stable state (Fig. 2e, f). However, this variability in the volatile-stable-volatile task was not strong, especially in the stable state, which suggests that participants perceived/expected the environment as having high uncertainty. A repeated measures ANOVA of the Order and Volatility showed different tendencies of estimated volatility between the stable-volatile-stable task and volatile-stable-volatile task *(F*_(1, 32)_ = 41.936, *p* < 0.001, *η* ^2^ = 0.567; Fig. S1). *Post hoc* analyses showed that estimated volatility differed as a function of an interaction with order; the volatile state (0.809) was significantly larger than the stable state (0.584) within the stable-volatile-stable task *(F*_(1, 32)_ = 40.85, *p* < 0.001), whereas the stable state (0.809) was significantly higher than the volatile state (0.712) within the volatile-stable-volatile task *(F*_(1, 32)_ = 7.65, *p* = 0.009).

We observed a decrease in learning when shifting from volatile to stable conditions across both ordered tasks (Fig. 2g). To confirm whether participants adapted their learning rates by order of transition (from stable to volatile state *vs*. from volatile to stable state), adapted learning rates were calculated by subtracting learning rate in the stable state from the volatile state. A repeated measures ANOVA of the Order and Transition showed that the adapted learning rates were significantly higher in volatile to stable transitions (adapted α_v-s_ = 0.076) than stable to volatile transitions (adapted α_s-v_ = −0.002), regardless of the task type *(F*_(1, 32)_ = 5.141, *p* = 0.03, *η* ^2^ = 0.138; Fig. 2h). Further analysis showed that adapted learning rate from volatile to stable state is significantly larger than zero which means that learning rate in the volatile state (0.4514) was larger than in the stable state (0.3714; *t*_(1, 33)_ = 2.413, *p* = 0.022). These results indicate that participants adapted their learning more in the transition from volatile to stable states than the other way around. The learning rate was not significantly different in the volatile environment from the stable environment *(p* > 0.05; Fig. S2). The adapted leaning rate in volatile to stable transitions was not significantly correlated to anxiety, depression or impulsivity *(ps* > 0.05; Fig. S3). Potential correlations were not explored for adapted learning rate in stable to volatile transitions due to lack of effect.

### Neuroimaging Results

The estimation of environmental volatility in the Bayesian model was negatively associated with activity in the bilateral caudate nuclei (left caudate, peak at x, y, z = −18, 12, 15, 87 voxels; right caudate, peak at x, y, z = 21, 15, 18, 54 voxels; *p* _*FWE*_ < 0.05; Fig. 3a). The increased blood oxygenation level-dependent (BOLD) signal in the bilateral caudate nuclei reflected lower estimated environmental volatility. Activity of the bilateral caudate nuclei in response to volatility was also associated with the variance of learning rates across the three blocks *(r*_(33)_ = 0.437, *p* = 0.01; Fig. 3b). These results revealed that encoding of environmental volatility in the caudate could predict the adjustment of learning strategies across the three blocks.

**Fig. 3.**
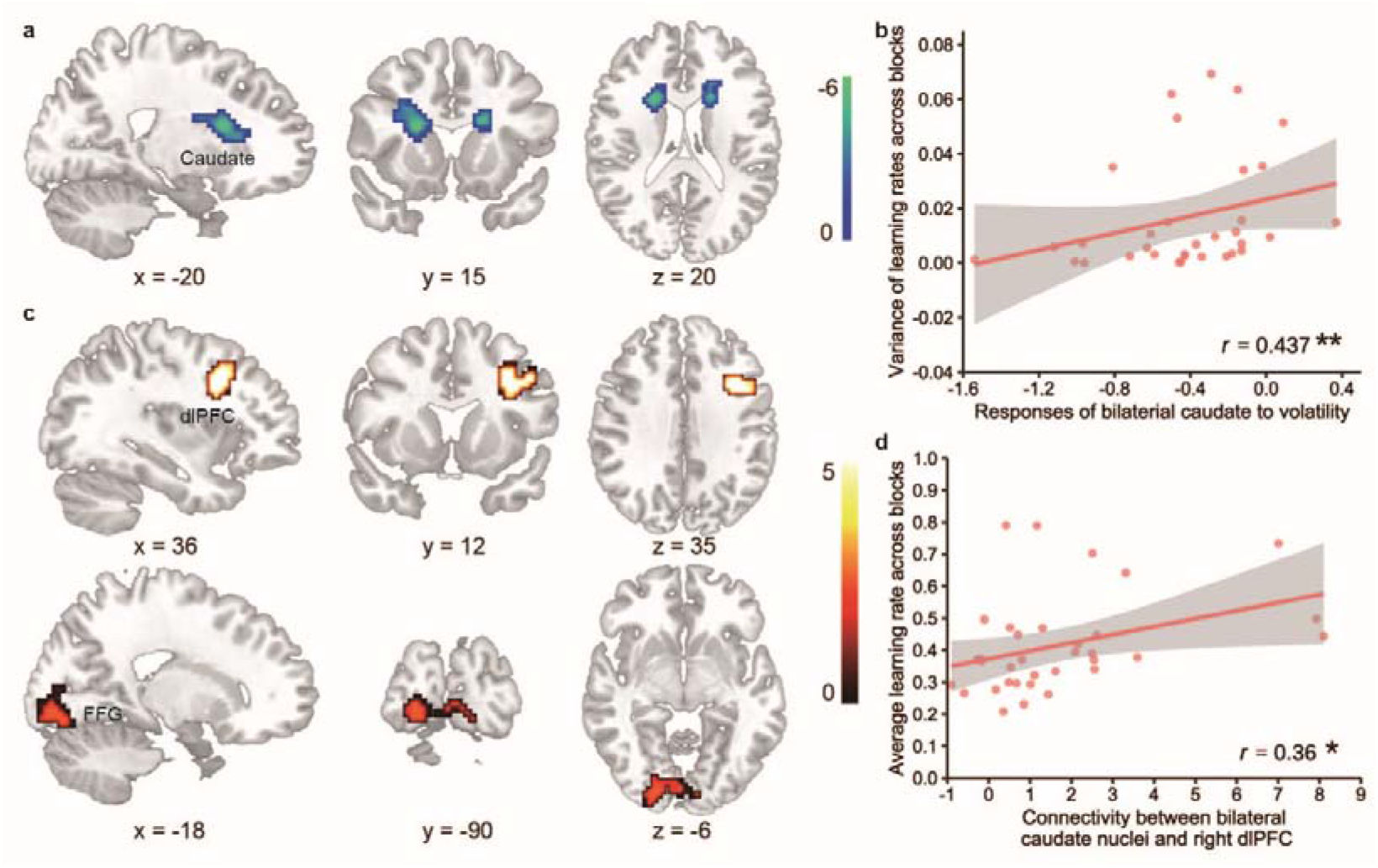
Brain activation. **a) Responses of bilateral caudate nuclei to estimated volatility.** Activity of the bilateral caudate nuclei (x = −18, y = 12, z = 15) was negatively associated with the estimated volatility in outcome evaluation. **b) Volatility related brain activity predicated changes in learning rates across blocks**. The degree to which bilateral caudate nuclei tracked estimated volatility could predict the variance of the learning rates across the three blocks. **c) Brain regions showed significant functional connectivity with bilateral caudate nuclei modulated by environmental volatility. d) Correlation between caudate-related connectivity and learning rate**.

To explore brain network configurations related to volatility, we performed PPI analysis, using the bilateral caudate nuclei as seeds. We found that the bilateral caudate nuclei showed positive connectivity with the right dorsolateral prefrontal cortex (dlPFC) and left fusiform gyrus (FFG) related to volatility (dlPFC, peak at x, y, z = −36, 12, 30, 93 voxels; FFG, peak at x, y, z = −18, −90, −6, 118 voxels; *p* _FWE_ < 0.05; Fig. 3c). Correlation analyses showed that functional connectivity between the bilateral caudate nuclei and right dlPFC was positively associated with averaged learning rate across the three blocks *(r*_(33)_ = 0.36, *p* < 0.05; Fig. 3d). Connectivity between the caudate and middle frontal gyrus (MFG) differed as a function of Order and Transition *(F*_(2, 64)_ = 3.587, *p* = 0.033, *η* ^2^ = 0.101; See Fig. 4b). *Post hoc* analysis revealed a significant decrease of caudate-MFG connectivity in the transition from volatile to stable relative to the stable to volatile transition, only for the stable-volatile-stable order *(F*_(1, 32)_ = 5.470, *p* < 0.05; Fig. 4c). To explore whether the caudate-MFG pathway tracked with volatility, we calculated correlations between environmental volatility and connectivity of the bilateral caudate and right MFG in each trial of each participant (Fig. S4). A one-sample *t*-test showed that correlations were not significantly different from zero *(ps*>0.05).

**Fig. 4.**
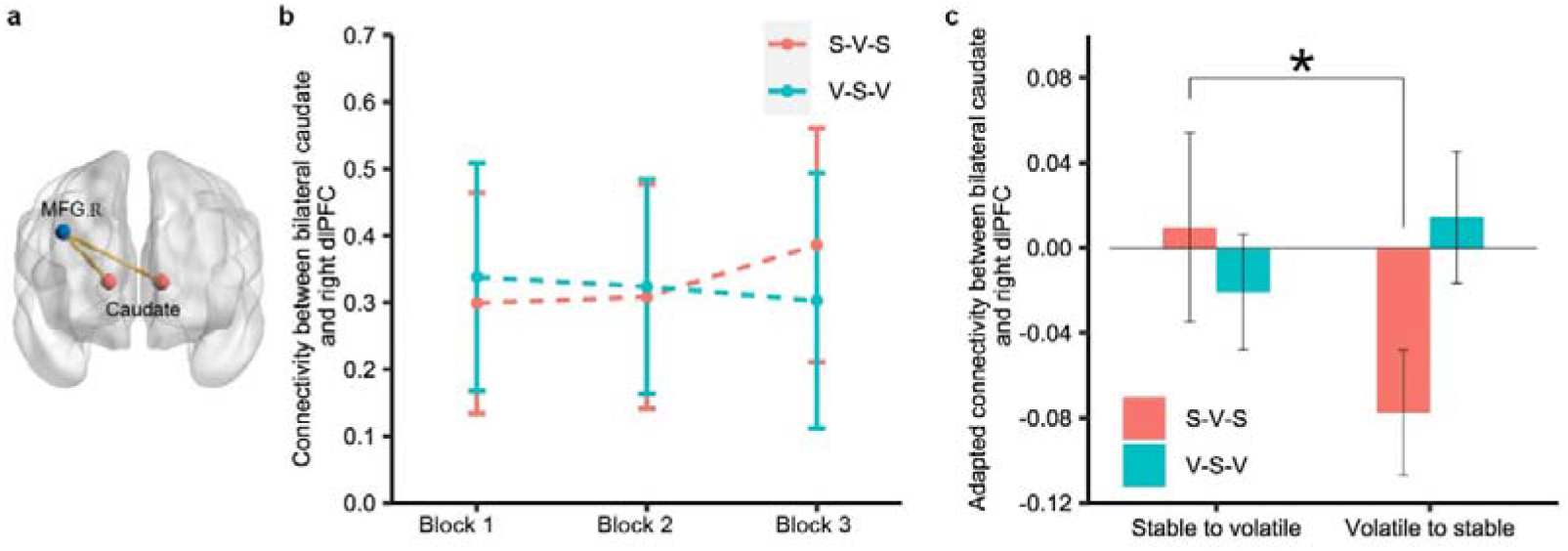
Dynamic functional connectivity. **a) The locations of regions of interest (the caudate) and its target region (the MFG).** Both of the two regions were defined based on the Anatomical Automatic Labeling (AAL) atlas). **b) The connection between the caudate and middle frontal gyrus across blocks. c) Changes of connectivity between the caudate and dlPFC between transitions from volatile to stable and from stable to volatile environment**.

Given that the brain dynamically adapts to the changing environment as a complex system, we examined dynamic modular structures of multilayer brain networks over multiple temporal scales. The modularity index was adapted to identify at which temporal scale the node was best partitioned into communities (Fig. 5a). Modularity analyses showed that *Q* at the smallest temporal scale (10s, one trial) was significantly higher than the other temporal scales *(ps* < 0.001; Fig. 5b). To examine changes of the community properties among the three blocks, we calculated the network flexibility within each block at 10s intervals (each block comprised 60 time-windows). Network flexibility decreased as a function of learning, especially from volatile to stable transitions (Fig. 5d). Repeated measures ANOVA of Order and Transition showed that adapted flexibility was significantly higher from volatile to stable transitions (adapted flexibility = 0.0054) than from stable to volatile transitions (adapted flexibility = −0.0019), regardless of the task type *(F*_(1, 32)_ = 4.672, *p* = 0.038, *η* ^2^ = 0.127; Fig. 5e). *Post hoc* analysis showed that adapted flexibility from volatile to stable transitions was not significantly larger than zero *(t*_(1, 33)_ = 1.589, *p* > 0.05). The contrast between volatile-to-stable and stable-to-volatile transitions showed that adapted flexibility was associated with activity in the left frontal pole (FP), right superior parietal lobule (SPL), left supramarginal gyrus (SMG), lateral occipital cortex, right OFC, left parahippocampal gyrus and brain stem *(ps* < 0.05; Fig. 5f). Correlation analyses didn’t show any significant relationships between adapted flexibility and adapted learning rates *(ps* > 0.05).

**Fig. 5.**
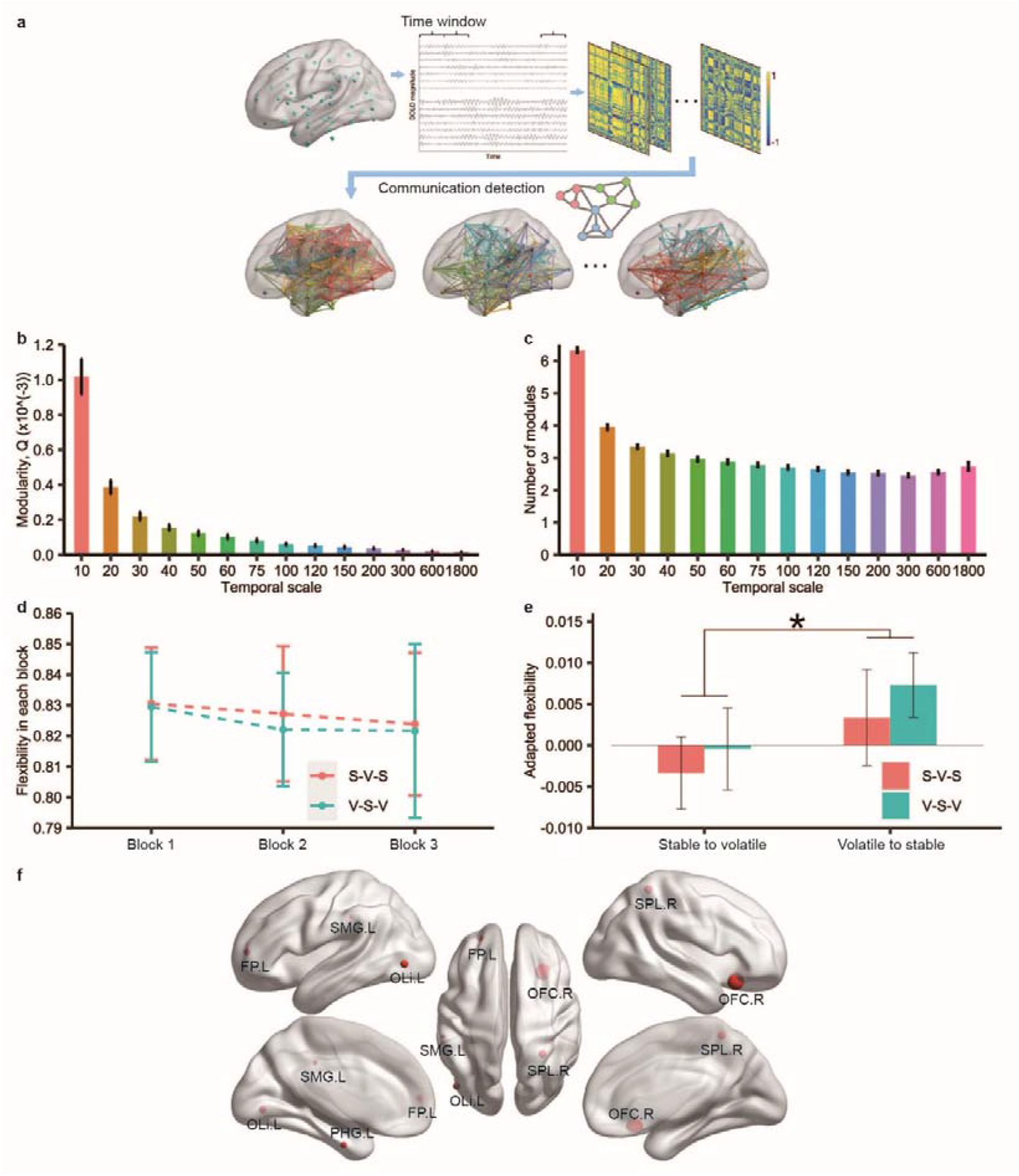
Dynamic brain networks. **a) Schematic overview of the dynamic brain network analysis.** The modular architectures of functional connectivity were detected by the modularity index. **b) Modularity index *Q* of community across several temporal scales. c) The number of modules of community across several temporal scales**. Error bars represent the standard error. **d) Network flexibilities in three blocks during the learning were calculated at 10s temporal scale**. Salmon dashed line for the stable-volatile-stable task; Dark turquoise dashed line for the volatile-stable-volatile task. Dots represent the mean of flexibilities. Error bars represent the standard deviation of network flexibilities in each block. **e) The adapted flexibility between transitions from volatile to stable and from stable to volatile environments)**. Error bars represent the standard error. **f) The brain regions represented adapted flexibility effect**. FP, frontal pole; SPL, superior parietal lobule; SMG, supramarginal gyrus; OLi, lateral occipital cortex; OFC, orbitofrontal cortex, PHG, parahippocampal gyrus; L, left; R, right.

## Discussion

The ability to adapt, on multiple levels, to constantly changing environments ensures optimal behavior for reward maximization and punishment minimization. Our results show distinctive behavioral adaptations and brain organizations in transitions between environments with varied volatility. Participants exhibited more adaptation and perceived higher volatility in stable environments among transitions from volatile to stable relative to stable to volatile environments. Subjective volatility of the environment was encoded in the bilateral caudate nuclei and its connectivity with the right dlPFC. Notably, modular organizations of dynamic caudate-dlPFC connectivity changed most within shorter time windows, reflecting the need for the brain to adapt quickly while learning of rapid environmental changes.

The asymmetrical adaptation between the volatile-stable and the stable-volatile transitions, suggest a greater need for adaptation under stable to volatile transitions. Heightened adaptation can be largely explained by early learning among major environmental changes. Participants exhibited greater adaptation under stable to volatile transitions when the first block was stable relative to when the first block was volatile. Asymmetrical adaptation is consistent with previous findings of transfer effects; shifts from a difficult task to an easier one is often more effortful than transfer in the opposite direction *(4)*. While previous studies have shown higher learning rates in the volatile environment and in the stable environment *(2, 11)*, our findings suggest important carryover effects such that experience with volatile environments may impact subsequent learning in a stable environment and vice versa. The idea has been supported in principle by a recent social norm violation study, which shows that participants preconditioned on unfair offers rejected comparable fair offers less frequently than participants preconditioned on generous offers *(21)*.

Several brain systems play key roles in adaptation to environmental transitions. The caudate nucleus, a critically part of the midbrain dopaminergic system, plays a key role in processing incentive salience *(22)*and learning associations between stimuli and responses *(17)*. A large number of studies have shown that the caudate encodes several learning parameters, including expected value *(23)*, action values *(24)*, reward prediction errors *(25, 26)*, and learning rates *(16)*. A primate study has shown that monkeys with caudate dopaminergic depletions exhibit marked difficulty in reconstructing the stimulus-reward associations after reversal *(15)*. The current findings show that the caudate also encodes estimated volatility, suggesting a new role of the caudate in learning. The present result is consistent with previous findings that the caudate is engaged in volatility estimates to fine-tune the weights of recent or remote prediction errors in predicting forthcoming needs for control *(27)*. Consistent with previous findings on the direct engagement of the dlPFC in encoding volatility *(9, 14)*, our results show a modulatory role of the connectivity between the caudate and right dlPFC in estimating volatility. Accumulating evidence has shown that the frontostriatal network is modulated by cognitive control *(28-30)*. Therefore, changes in the frontostriatal pathway associated with cognitive control might be involved in estimates of environmental volatility. Contrary to previous work *(1, 2, 9, 31)*, we did not find robust volatility-related signals in the ACC. One potential explanation might be that the magnitude of outcome in our task was fixed rather than variable as in previous studies *(2, 11)*. Changing outcome probabilities and outcome magnitudes may have increased task difficulty in prior work. It is possible that strong contrast effects (e.g. high probability with low magnitude vs. low probability with high magnitude) might drive ACC activity in previous studies.

In transition from the volatile-stable direction to the stable-volatile direction, participants showed higher network flexibility, suggesting a dynamic reorganization of the brain in adaptation to environmental volatility. Dynamics of functional brain connectivity has been widely used to examine the complex human cognition of moment-by-moment changes in learning *(32-34)*, by providing dynamic measurements of flexibility in coordination among different brain states in responses to adaptative behaviors *(35, 36)*. Based on dynamic functional connectivity, modular structures aggregated by small subsystems or modules might facilitate behavioral adaptation *(19)*. These findings point to a pivotal role of the frontoparietal control network in the adapted flexibility of individuals in transitions between environments with different volatility.

In conclusion, to the best of our knowledge, the current work is the first to examine the way that the human brain adapts to transitions in environmental volatility. Our results show asymmetrical behavioral and neural adaptations during the environmental transitions with superior adaptation under transitions from stable to volatile environments rather than the opposite. These flexible adaptations are modulated by the striatal reward system and its dynamic connections with the prefrontal control system. The current work sheds light on the nature of the human brain adaptations to navigation of variable environments in daily life.

## Materials and Methods

### Participants

Thirty-Seven (20 females) Chinese participants, aged between 18 and 30 years (mean± SD = 21.62± 2.79 years), without any history of psychiatric disorders were recruited from several Universities in Beijing. Three participants were excluded from all analyses as they missed more than 10 of the 180 trials during the primary task (5.6%). The study protocol was approved by the local Ethics Committee. All participants provided informed consent.

### General procedure

To ensure participants understood the probability and reversal components underlying the task, a three-stage training task was implemented *(37)*. Participants then completed an adjusted probabilistic reward reversal learning task while undergoing fMRI. Participants completed the trait subscale of the Spielberger’s State-Trait Anxiety Inventory *(38)*, the Self-rating Depression scale *(39)* and Barrett impulsivity scale *(40)*. After the experiment, all participants were fully debriefed and received payment based on their task performance.

### Task design

A probabilistic reward reversal learning task (Fig. 1a) was adapted from previous reversal learning tasks used to examine learning strategies in environmental volatility *(2, 11)*. The task consists of three blocks of 60 trials in which participants were required to make choices between two options with specific reward probabilities (Fig. 1b, 1c). In the stable block, two options were stably associated with reward probabilities at 75% and 25%, respectively. In the volatile block, the probabilities of the two options switched between high (80%) and low (20%) reward probability every 20 trials. To test whether adaptations differ between the transition from stable to volatile and the transition from volatile to stable blocks, participants were randomly assigned to complete the three blocks either in the order of stable-volatile-stable or volatile-stable-volatile without taking a break. None of the participants was informed that tasks would be divided into three different blocks.

In each trial, two options with horizontal and vertical gratings were presented with a visual angle of approximately 8º . Participants were required to make decisions within 3s. Once they responded, the selected option would be highlighted via a white frame to acknowledge the choice with a duration of 0.2s. Sequentially, a question mark was presented at the center of the screen with a jitter interval at 2-4s, to indicate that the outcome was pending. At the phase of outcome, reward (green “+ 1”) or no reward (red “+ 0”) was presented at the center of the screen for two seconds. At the end of trial, an inter-trial interval with a fixation cross was presented for 0.8-5.8s, to ensure total trial duration was 10s.

### The Rescorla-Wagner model

Participants’ learning rates in three blocks were estimated by a simple Rescorla-Wagner (RW) learning model *(41)*. In (*i* + 1)^*th*^ trial, the predicted reward probability of option A, 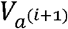, was updated using the following equation,

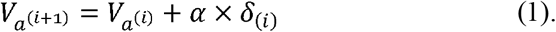

The 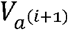 and 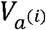 represent the expected values. Specifically, the predicted reward probability of the chosen option A for the (*i* + 1)^*th*^ and *i*^*th*^ trial. The *α* ∈ [0, 1] indicates the learning rate and *δ*_*(i)*_ is the prediction error on the *i*^*th*^ trial which was calculated by comparing the actual reward *r*_*(i)*_ with the expected value 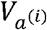,

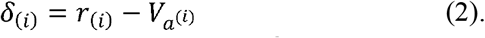

The likelihood of the option chosen by participants on the *i*^*th*^ trial was estimated by a sigmoidal probability distribution,

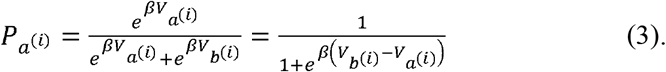

Here, *β* ∈ [0, 10]indicates the inverse temperature parameter which controls the degree to which an option would be chosen. When 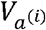 is larger than 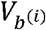, the larger the *β*, the higher probability that option A would be chosen.

Participants were instructed that the difference between the probabilities of two options would be clear without mentioning that they were opposite, and both add to one. Although it is theoretically possible for participants to learn about the probabilities of two options independently, the probability of the unchosen option B was calculated as follows,

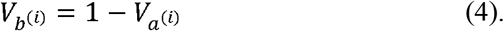

The probability of the second option was fixed to make learning simpler.

### The Bayesian learner model

We also assume that participants would track the probabilities of option and outcome optimally by a Bayesian rule which has been described in previous studies *(1, 2, 11)*. Here, we provide a brief overview. In each trial, the outcome *y*_*(i)*_ reward or not, was determined by the underlying estimated probability of option *r*_*(i)*_:

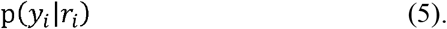

The probability of option in the *i*^*th*^ trial, *r*_*(i)*,_ was determined by the probability of the option on the (*i* − 1)^*th*^, *r*_*(i −* 1),_and the estimated volatility on the *i*^*th*^, *ν*_*(i)*_:

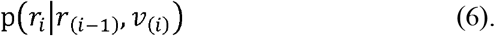

The volatility *v* refers to an estimate of the expected rate of change of *r*. According to the Bayesian model, a low volatility would be estimated and each new outcome would have little influence on the estimate of *r* in a stable environment, whereas a high volatility would be estimated in a fast-changing environment, implying *r* may be expected to change quickly. The model also assumes that the way participants track environmental volatility is same as the way participants track the changing probability,

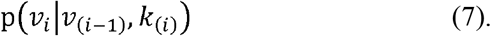

The environmental volatility on the *i*^*th*^, *ν*, was determined by the volatility of the preceding trial, *ν*_*(i −* 1)_and the control parameter, *k*_*(i)*._ A large k implies that stable or volatile environment switches to the other one frequently.

All parameters representing the participant’ expectations of the statistics of the environment were modeled by the joint probability distribution,

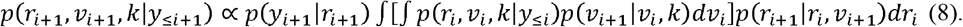

### Model fitting

The learning rates and inverse temperature parameters were firstly fitted by a grid search. However, while increasing the precision of two parameters (e.g., *α* [0:0.01:1] and *β* (0:0.1:10]), the number of all possible combinations of two parameters across three blocks via grid search requires a very large amount of memory. To resolve this issue, we first adopted the solution that these parameters would be estimated separately for each of the three blocks by the grid search. The first prediction values in block 2 and block 3 (trial 61 and 121) were from previous estimates (trial 60 and 120). We also adopted the function fMINSEARCH (fMIN) in Matlab to estimate the parameters without having to measure every single point. To determine the best-fitting values of the learning rates and inverse temperature parameters, we used both the maximum likelihood estimate (ML) and expected value estimate (EV) (multiplying the value of each bin with the probability of that value). Additionally, trials with no response and trials in which reaction time less than 200ms were excluded.

### Model selection

The Bayesian information criterion (BIC) was used to assess which model best captured participants’ choices. Lower value indicates the better fitting. *L* is the likelihood, *m* is the number of estimated parameters, *N* is the number of observations. The Bayesian observer model includes 3 parameters *(r, v, k)*. The RW model with fMINSEARCH that can look for minima without every single point includes 6 parameters (*α, β* per block). The RW model with Grid search includes 2 parameters. Since this model fitted the participants’ choices separately for three blocks, the BIC of each block was first calculated and then added together to get the total BIC for each participant.

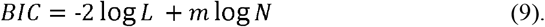

To examine the fitting effect of model, we calculated the correlation of the participants’ choices with probabilities estimated by the RW and Bayesian model. The average correlation coefficient was then used as a corroboration to indicate which model was best.

### fMRI data acquisition

MRI data were collected on a 3T Siemens Prisma MRI scanner with a 64-channel head-neck coil at the Center for MRI Research, Peking University. Functional MRI images were acquired with a simultaneous multiband echo planar imaging (EPI) sequence (TR/TE = 1000/30 ms; FOV = 224 × 224 mm; matrix = 64 × 64; slice thickness = 3.5 mm; slice number = 34; flip angle = 73°; multiband factor = 2). High spatial resolution T1-weighted anatomical images were obtained with the magnetization □prepared rapid gradient□echo (MPRAGE) sequence (TR/TE = 2530/2.98 ms; FOV = 256 × 256 mm; matrix = 256 × 256; slice thickness = 1 mm; flip angle = 7°).

### fMRI data preprocess

All images were preprocessed using SPM12 (Wellcome Trust Center for Neuroimaging, http://www.fil.ion.ucl.ac.uk/spm/software/spm12/). The fMRI data were first corrected for head motion, and the realigned images were coregistered to the T1 structural image which had been manual reoriented to the anterior commissure and then segmented into white matter, gray matter, cerebrospinal fluid (CSF), bone, soft tissues, and air using default tissue probability maps of SPM12. The images were then normalized to the standard Montreal Neurological Institute (MNI) space with final resolution of 3 mm^3^. Finally, normalized images were smoothed with a Gaussian kernel of 6 mm full-width at half-maximum.

### fMRI data analysis

To identify the brain responses to the environmental volatility, we constructed a general linear model (GLM) with the onsets of the options presentation modulated by a parametric regressor (the environmental volatility), onsets of the choice selection, onsets of the jitter period modulated by the volatility, onsets of the feedback delivery modulated by the volatility, onsets of trials with insufficient response time (< 200 ms, including non-response trials), onsets of trials with large head motion (the framewise displacement, FD > 0.5). Additionally, six estimated head movement regressors and a FD regressor were also included as covariates of no interest. The regressors in the GLM design matrix were then convolved with the canonical hemodynamic response function (HRF). Group analyses for brain activation were performed with a random-effect model using a one-sample *t*-test. All results were whole-brain corrected for multiple comparison by a voxel-wise uncorrected threshold at *p* < 0.001 with a family-wise error (FWE) corrected for cluster-level at *p* < 0.05.

### Psychophysiological interaction analyses

To identify the volatility-specific changes in the interaction between brain regions in whole brain functional connectivity, the psychophysiological interaction (PPI) was performed by using the significant cluster as seed region of interest (ROI) which was related with estimated volatility. For each participant, the first eigenvariate of the ROI was extracted to get the individual voxel time-course. In order to generate the PPI interaction term, this time-course was deconvolved with the canonical HRF and then multiplied by the vector of the estimated volatility. This interaction term was then convolved with the canonical HRF and entered into a PPI GLM along with the vectors of the onsets for the estimated volatility during the feedback, the original eigenvariate time-course and covariates of no interest (six head movement and a FD). After that, we performed second-level analyses with one-sample *t*-test for the contrast images of the PPI interaction term. The results were corrected for a voxel-wise uncorrected thresholded at *p* < 0.001 with an extent FWE-corrected cluster-level *p* < 0.05.

### Dynamic Functional connectivity

To examine dynamic changes between the seed-ROI and the significant clusters from PPI analysis, we extracted BOLD signals from the corresponding ROIs based on the AAL atlas. The linear Pearson’s correlations between the seed and target regions were calculated in three blocks to estimate individual dynamic functional connectivity.

### Dynamic Brain Networks

To explore modular organizations of the brain in the dynamic adaptation, the modular structures spanning several temporal scales were constructed according to previous recommendations *(19)*. The smoothed images were temporally detrended to reduce the effects of linear drift and nuisance signals were removed to reduce the effects of non-neuronal fluctuation, including head motion, the white matter and the CSF. fMRI data were then band-pass filtered to reduce the effects of low frequency drift and high-frequency physiological noises with 0.06-0.12 Hz *(19)*. The whole brain was parcellated into 112 ROIs identified in the 3mm Harvard-Oxford (HO) atlas. To construct the individual functional connectivity matrices, the mean BOLD time series were first estimated by averaging voxel time series in each ROI and the linear Pearson’s correlation *r*_*ij*_ between all pairs of ROIs *i* and *j*. To correct for multiple comparisons, we first computed the *p*-values *p*_*ij*_ of each *r*_*ij*_ using the MATLAB function *corrcoef* and then tested the significance of *p*_*ij*_ using a False Discovery Rate (FDR) of *p* < 0.05. The *p*_*ij*_ of the correlation matrix elements *r*_*ij*_ which passed the FDR corrections was retained. Otherwise, the correlation matrix elements *r*_*ij*_ were set to zero. These corrected matrices 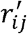 constituted adjacency matrices **A**, elements of which 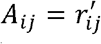. Based on our experimental setting, we measured functional connectivity over several temporal windows: [10s, 20s 30s, 40s, 50s, 60s, 75s, 100s, 120s, 150s, 200s, 300s, 600s, 1800s]. The node in each functional connectivity matrix was partitioned into a community by maximizing the modularity index *Q (42)*. To identify organizations of subtle networks, we constructed the undirected weighted graphs that preserving the information of the strength of connections *r*_*ij*_ *(43)*. A spectral optimization algorithm was used to optimize the modularity index *Q (44)*. The most popular formula of *Q* was used *(44)* as follows,

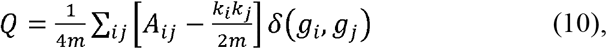

where 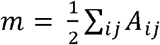, *k*_*i*_ is the strength of node, *i, k*_*j*_ is the strength of node *j*. When node *i* and node *j* are in the same module, *δ (g*_*i*_, *g*_*j*_) = 1; Otherwise, it equals 0. Both positive and negative weighted correlation coefficients were used to construct the community with the assumption that it would provide more useful information about the modularity partitions than only using positive correlation matrix elements *(45)*. The modularity was generalized *(46)* as

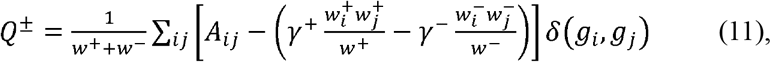

where *γ*^+^ and *γ*^−^ are resolution parameters which usually set as one for simplicity and 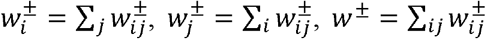. Although it has been argued that *Q*^+^ and *Q*^−^ should be treated equally because positive and negative connections play different roles in functional brain networks *(45)*, we defined an asymmetric formula of modularity as below based on previous studies which proposed that high-*Q*^+^modularity partitions are more optimal than high-*Q*^−^objectively *(45)*,

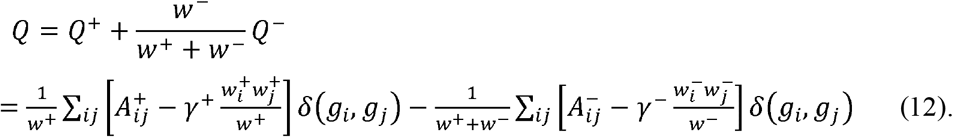

The maximization of the modularity index *Q* categorizes the nodes into the communities such that the total edge weight within the module is as large as possible. Hence, we selected one temporal window with the largest *Q* over the several temporal windows. To measure changes in the modules during the learning, we calculated the *flexibility* of a node *f*_*i*_ at each block. The *flexibility* was defined as the number of times the node changed modular assignment throughout the block, normalized by the number of all possible changes*(19)*. The flexibility of the network in each block was calculated as below,

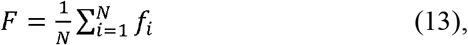

where time window *N = 600/temporal scale*.

## Supporting information

manuscript

## Funding

National Natural Science Foundation of China (31920103009)

National Natural Science Foundation of China (31871137)

Major Project of National Social Science Foundation (20&ZD153)

Young Elite Scientists Sponsorship Program by China Association for Science and Technology (YESS20180158),

Guangdong International Scientific Collaboration Project (2019A050510048)

Guangdong Key Basic Research Grant (2018B030332001)

Shenzhen-Hong Kong Institute of Brain Science-Shenzhen Fundamental Research Institutions (2019SHIBS0003)

Shenzhen Science and Technology Research Funding Program (JCYJ20180507183500566 and JCYJ20180305124819889)

## Author contributions

Conceptualization: P.X., H.A

Methodology: P.X., J.X

Investigation: J.X

Supervision: P.X., H.A

Writing—original draft: J.X., N.T.-D., Y.L., A.A., H.A., P.X

Writing—review & editing: J.X., N.T.-D., Y.L., A.A., H.A., P.X

## Data availability

The data that support the findings of the current study are available from the corresponding author (P. X.) upon request.

## Code availability

The custom scripts used to analyze are available from the corresponding author (P. X.) upon request.

