## Supplementary material for "Asymmetrical adaptations to increases and decreases in environmental volatility": manuscript

*Corresponding author:

Hui, Ai, Ph.D.

Supplementary Text

Supplemental Methods

Training task. The training task was adapted from a previous probabilistic reversal learning task (*1*). This training program consisted of three stages from easy to difficult. The first stage was a simple reversal task, in which option A resulted in a reward 100% and option B resulted in a reward 0%. After four consecutive choices of the correct option, a reversal would occur with 50% probability. The first training stage was ended after the three reversals. The second training phase was simple probabilistic task, in which option A resulted in a reward 70% and option B resulted in a reward 30% as in the experiment but the probability did not reverse. The second stage was ended after the six consecutive correct choices. The third training stage combined the probability with reversal. After four consecutive choices of the correct option with 70% probability of reward, a reversal would occur with 50% probability. The final training stage was ended after the two reversals. Same gratings stimuli were used in the training stages with those used in the experiment. Participants were informed that their remuneration was not depended on their performance during the training stages.

Supplemental Results

Behavioral Results. The sum of the rewards (i.e., scores) across the blocks that individuals received in the volatile environment was significantly higher than in the stable environment (F(1, 32) = 5.507, p = 0.025, η2 = 0.147). It is worth noting, however, that participants may perform better in the volatile (relative to the stable), simply because scores in the volatile environment were affected by noise (Fig. S5c). We found no significant differences between the stable and volatile environment in accuracy and reaction time (Fig. S5a, b). These results suggest that difficulty may be equivalent in the stable and volatile environments as the true reward likelihood was slightly higher in the volatile (0.8) than in the stable (0.75) environments, consistent with the previous study (Behrens et al., 2007). There were also no significant differences between the stable-volatile-stable task and the volatile-stable-volatile task in accuracy, reaction time or score (p > 0.05) (Fig. S5d, e, f).

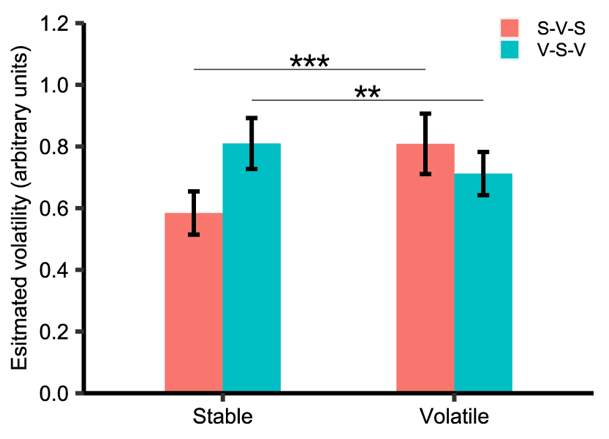

Fig. S1. Estimated volatility in the stable and volatile state for stable-volatile-stable task and volatile-stable-volatile task.

There was a significant interaction effect between the Task and State on the estimated volatility (F(1, 32) = 41.936, p < 0.001, η2 = 0.567). Further analysis showed that estimated volatility in the volatile state (0.809) is significantly larger than it in the stable state (0.584) in the stable-volatile-stable task (F(1, 32) = 40.85, p < 0.001) whereas, estimated volatility in the volatile state (0.712) is significantly less than it in the stable state (0.809) in the volatile-stable-volatile task (F(1, 32) = 7.65, p = 0.009).

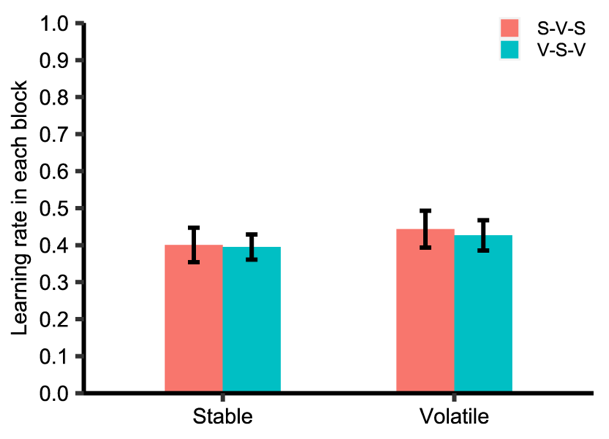

Fig. S2. Learning rate in the stable and volatile state for stable-volatile-stable task and volatile-stable-volatile task.

There was no significant difference between the stable and volatile state in learning rate between two tasks (p > 0.05).

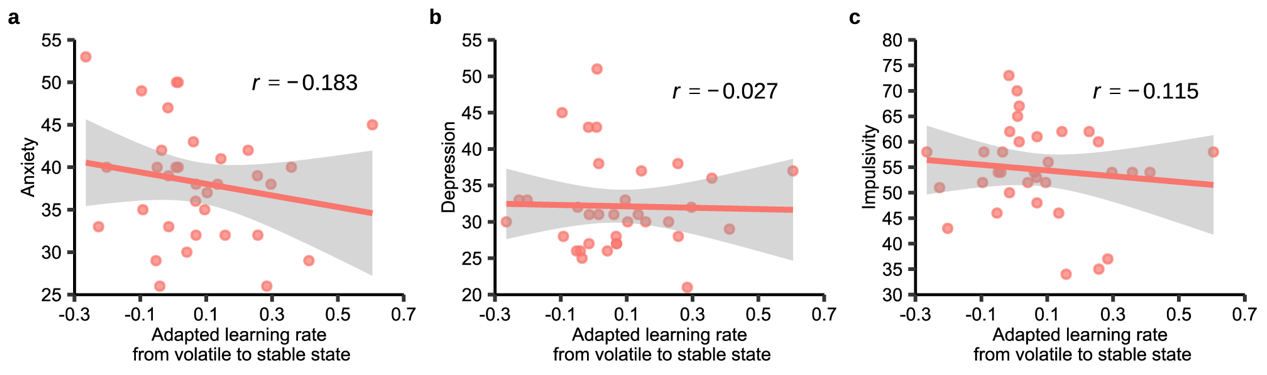

Fig. S3. Relationship between adapted learning rate and trait anxiety, depression and impulsivity.

The adapted leaning rate from volatile to stable state was not significantly correlate to the anxiety, depression or impulsivity (*ps* > 0.05).

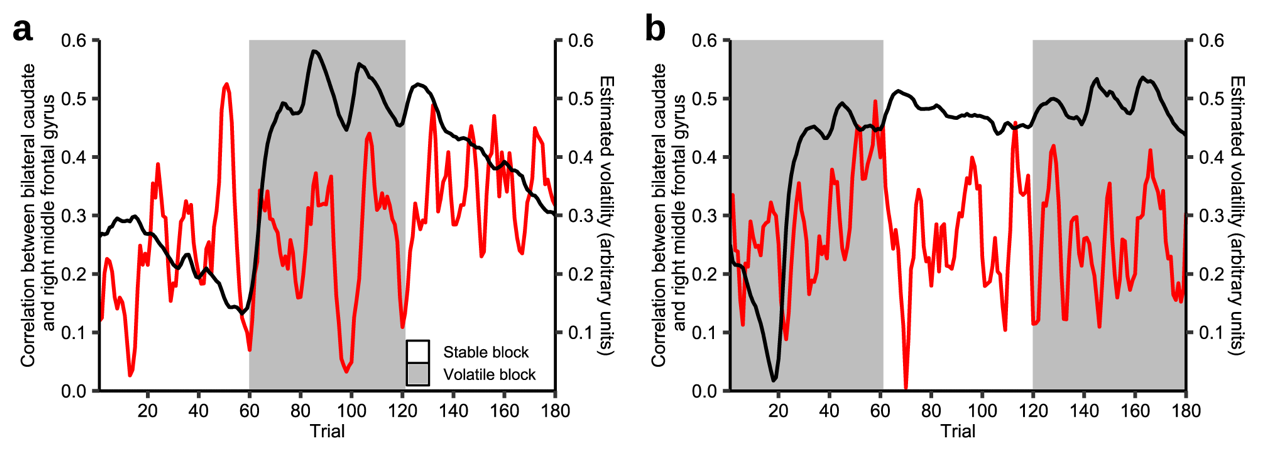

Fig. S4. Connection between caudate and middle frontal gyrus and estimated volatility across the whole experiment.

**a)** **Stable-Volatile-Stable task**. Individual functional connectivity between the bilateral caudate and right MFG was calculated at trial-level. Then, the correlation between estimated volatility and caudate-MFG connection was calculated at subject-level. One-sample *T* test showed that the correlation was not significantly different from zero for stable-first group (*p* > 0.05). Mean caudate-MFG connection and estimated volatility were displayed for stable-volatile-stable task. **b)** **Volatile-Stable-Volatile task**. One-sample *T* test showed that the correlation was not significantly different from zero for volatile-first group (*p* > 0.05). Mean caudate-MFG connection and estimated volatility were displayed for volatile-stable-volatile task.

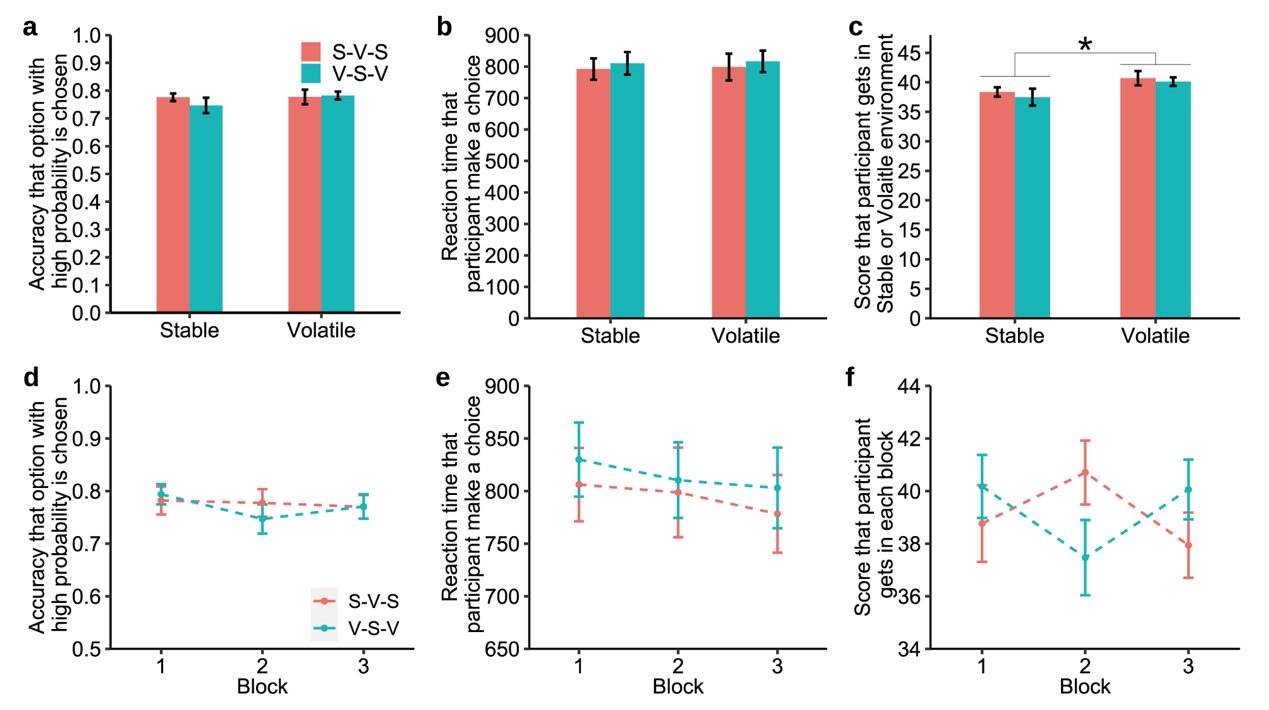

Fig. S5. Behavioral results.

**a) Accuracy in the stable and volatile state for stable-volatile-stable task and volatile- stable-volatile task.** There was no significant difference between stable and volatile state in accuracy between two tasks (*p* > 0.05). **b) Reaction time in the stable and volatile environment for stable-volatile-stable task and volatile-stable-volatile task.** There was no significant difference between stable and volatile state in reaction time between two tasks (*p* > 0.05). **c) Score in the stable and volatile state for stable-volatile-stable task and volatile-stable-volatile task.** The score in volatile state was significantly larger than it in stable state (*F*_(1, 32)_ = 5.507, *p* = 0.025, *η*^2^ = 0.147). **d) Accuracy across three blocks.** There was no significant difference between the stable-volatile-stable task and volatile- stable-volatile task in accuracy across three blocks (*p* > 0.05). **e) Reaction time across three blocks.** There was no significant difference between the stable-volatile-stable task and volatile- stable-volatile task in reaction time across three blocks (*p* > 0.05). **f) Score across three blocks.** There was no significant difference between the stable-volatile-stable task and volatile- stable-volatile task in score across three blocks (*p* > 0.05).
